# Integrative Genomic and Structure-Based Prioritization of Molecular Targets in Multidrug-Resistant *Salmonella enterica* from Brazilian Poultry

**DOI:** 10.64898/2026.05.17.723367

**Authors:** Juan Philippe Teixeira, Daniel Ferreira de Lima Neto, Chiara Brancalion, Miklos Maximiliano Bajay, Pedro Filipe de Souza Teles, Rafael Straus de Sá, Saad Khan, Thales Quedi Furian, Lenita de Cássia Moura Stefani

**Affiliations:** State University of Santa Catarina – Center for Agricultural and Veterinary Sciences – Graduate Program in Biochemistry and Molecular Biology (PMBqBM), Lages, SC, Brazil; State University of Santa Catarina – Graduate Program in Animal Science (PPGZOO), Chapecó, SC, Brazil; Federal University of Rio Grande do Sul – Center for Diagnosis and Research in Avian Pathology (CDPA), Porto Alegre, RS, Brazil

**Keywords:** *Salmonella*, antimicrobial resistance, whole-genome sequencing, molecular docking, poultry production

## Abstract

*Salmonella* spp. remains one of the leading foodborne pathogens worldwide, and the circulation of multidrug-resistant strains in the poultry industry poses a significant challenge. In this study, five isolates from poultry litter swabs (commercial broiler chickens) belonging to the *Salmonella* Heidelberg and *Salmonella* Minnesota serovars were characterized using an integrated approach involving phenotypic resistance profiling, whole-genome sequencing, structural prioritization of molecular targets, and *in silico* screening of ligands. All isolates exhibited multidrug resistance phenotypes and genetic repertoires consistent with resistance to β-lactams, sulfonamides, and tetracyclines, as well as determinants linked to efflux systems, virulence, and persistence. Genomic analysis allowed for the prioritization of five proteins for structural investigation: CTX-M-2, CMY-2, Sul2, AcrB, and SpvC. Sequence-structure validation revealed high correspondence between the proteins of the isolates and the experimental structures selected for CMY-2, Sul2, AcrB, and SpvC, while CTX-M-2 was modeled with high structural confidence. Molecular docking analyses with GNINA revealed distinct behaviors among the targets. Sul2 showed biological relevance but a more conservative structural response, with no significant gain after analog generation. In contrast, AcrB stood out as the most promising target, with analogs generated by BRICS yielding better scores and, in some cases, coherent international networks identified by PLIP. The results demonstrate that the integration of phenotype, comparative genomics, and structural prioritization constitutes a rational strategy for selecting targets and molecular candidates in multidrug-resistant avian strains of *S.* Heidelberg and *S.* Minnesota.

## INTRODUCTION

Nontyphoidal salmonellosis remains one of the leading foodborne bacterial zoonoses worldwide, causing gastroenteritis associated with the consumption of contaminated products, especially those of animal origin. Transmission frequently occurs through foods such as eggs, meat, and poultry products, making surveillance of this pathogen particularly important in intensive poultry production systems [1].

In Brazil, the poultry industry is of great economic and public health importance, and recent genomic studies show that certain serovars of Salmonella enterica are playing a prominent role in this sector. Among them, Salmonella Heidelberg and S. Minnesota have been described as prevalent or emerging serovars in Brazilian poultry production, with circulation also detected in exported products, reinforcing their potential impact on public health and international trade [2].

This situation is exacerbated by the spread of antimicrobial resistance. In Brazilian isolates of S. Heidelberg and S. Minnesota from poultry, resistance profiles to third-generation cephalosporins and fluoroquinolones have already been described, in addition to the presence of relevant genes such as blaCMY-2, associated with plasmid replicons, and sets of determinants linked to multidrug resistance and virulence [3]. In a more recent study of S. Heidelberg isolated from broiler chickens in Brazil, a high frequency of antimicrobial resistance and association with multiple resistance and virulence genes was observed, reinforcing the epidemiological potential of this serovar in the poultry industry [4].

Similarly, S. Minnesota has been recognized as an emerging serovar in Brazilian poultry farming. Recent data obtained from isolates derived from chicken litter in southern Brazil showed that samples of this serovar exhibited a multidrug-resistant profile confirmed both phenotypically and genetically, which supports concerns regarding its persistence and spread in the production environment [5]. Furthermore, analyses of environmental persistence and genotypic/phenotypic profiles indicate that S. Minnesota is frequently associated with the poultry production chain and with genes such as tetA and blaCMY-2, reinforcing its adaptation to this niche [6].

As a result, whole-genome sequencing (WGS) has become a key tool for the surveillance of foodborne pathogens. This approach allows for high-resolution characterization of the resistome, virulome, plasmidome, and genetic relationships among isolates, expanding the capacity for epidemiological tracking and interpretation of the mechanisms involved in antimicrobial resistance [7]. In Salmonella surveillance, the use of WGS has also been adopted by food safety authorities and monitoring networks, precisely because of its value for typing, outbreak investigation, and genomic comparison among isolates from different sources [8]

However, the genomic identification of resistance and virulence genes, while essential, is not sufficient on its own to indicate which molecular targets hold the greatest potential for functional and pharmacological exploitation. Proteins such as β-lactamases, enzymes associated with sulfonamide resistance, and components of multidrug efflux systems constitute particularly relevant targets because they represent central mechanisms of bacterial survival under antimicrobial pressure. In the case of efflux pumps, there is recent evidence that systems such as AcrAB/AcrD in Salmonella remain important targets in in silico and experimental studies aimed at discovering inhibitors [9]. More broadly, efflux pumps in Gram-negative bacteria are recognized as key components of multidrug resistance and, therefore, as promising targets for chemotherapeutic interventions [10].

Thus, approaches that integrate resistance phenotypes, comparative genomics, and structural prioritization through molecular modeling and docking can offer a rational strategy for transforming surveillance data into more refined mechanistic hypotheses. This integration allows for the reduction of redundancies, the selection of targets biologically consistent with the profile observed in the isolates, and, at the same time, the guidance of the initial screening of ligands with the potential to interact with proteins associated with resistance and virulence. The overall analytical design is summarized in Figure 1. In light of this overview, the present study sought to characterize multidrug-resistant isolates of S. Heidelberg and S. Minnesota from the Brazilian poultry industry, integrating phenotypic, genomic, and structural data for the rational selection of molecular targets of interest.

**Figure 1.**
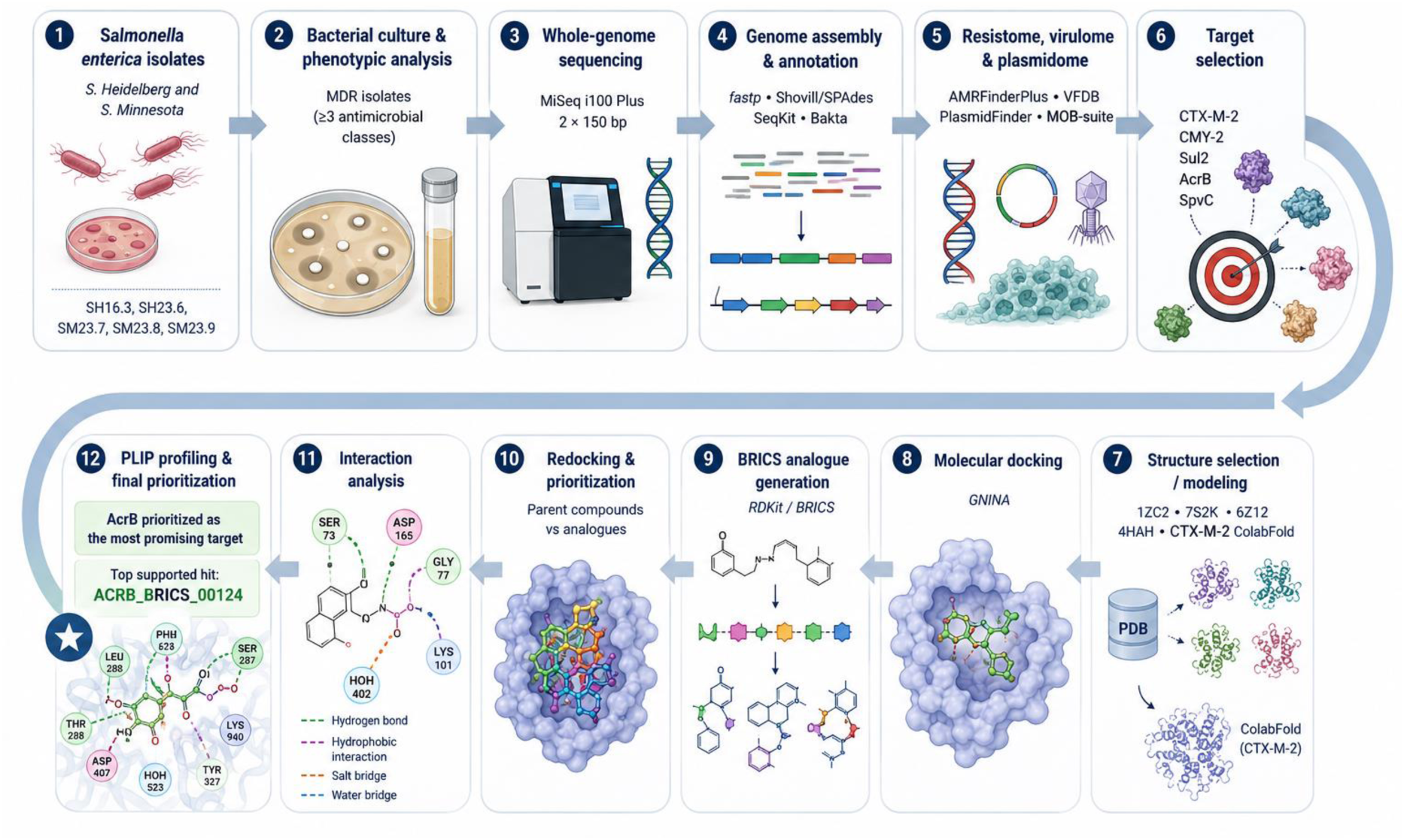
Integrated workflow for genomic and structure-based prioritization of molecular targets. The workflow starts with the selection of multidrug-resistant Salmonella enterica isolates from poultry, followed by phenotypic resistance profiling, whole-genome sequencing, assembly, annotation, resistome/virulome screening, structural target selection, molecular docking with GNINA, BRICS-based analog generation, PLIP interaction profiling, and final prioritization of candidate complexes.

## MATERIALS AND METHODS

### Selection of strains, origin of isolates, phenotypic profile, and criteria for multidrug resistance

The Salmonella enterica strains used in this study were serotyped and provided by the Center for Diagnosis and Research in Avian Pathology (CDPA) at the Federal University of Rio Grande do Sul (UFRGS). The isolates originated from swabs taken from companies in the broiler chicken production chain in Brazil, representing strains of sanitary and economic importance to the national poultry industry. For the present study, five strains were selected based on their phenotypic, genotypic, and structural relevance: SH16.3, SH23.6, SM23.7, SM23.8, and SM23.9. All included strains were previously classified as multidrug-resistant based on phenotypic antimicrobial susceptibility testing (antibiogram), as they exhibited resistance to three or more classes of antimicrobials. In this study, phenotypic resistance data were used as a biological eligibility criterion for including the strains in the genomic pipeline and, subsequently, as an interpretive reference for prioritizing molecular targets in the in silico stage. Among the five selected strains, distinct resistance profiles were observed. Strain SH16.3, belonging to the S. Heidelberg serovar and isolated in 2016 in the state of Santa Catarina, exhibited resistance to gentamicin, enrofloxacin, nalidixic acid, and tetracycline. Strain SH23.6, also belonging to the S. Heidelberg serovar and isolated in 2023 in Rio Grande do Sul, showed resistance to colistin, amoxicillin/clavulanic acid, and enrofloxacin. Among the S. Minnesota isolates from 2023, strain SM23.7 exhibited resistance to ampicillin, chloramphenicol, cefotaxime, tetracycline, enrofloxacin, and sulfamethoxazole-trimethoprim; strain SM23.8 exhibited resistance to ampicillin, tetracycline, enrofloxacin, and sulfamethoxazole-trimethoprim, as well as additional resistance to amoxicillin/clavulanic acid, enrofloxacin, and ceftiofur in a supplementary assay; and strain SM23.9 showed resistance to ampicillin, aztreonam, cefotaxime, tetracycline, azithromycin, and sulfamethoxazole-trimethoprim, as well as additional resistance to amoxicillin/clavulanic acid in a supplementary assay. This diversity of phenotypic profiles, combined with the geographical origin and biological representativeness of the isolates, supported the selection of these five strains as the core of the genomic, structural, and cheminformatics analyses conducted in this study.

### Bacterial reactivation, culture preparation, and genomic DNA extraction

Brain Heart Infusion (BHI), Rappaport-Vassiliadis, and XLD (Xylose-Lysine-Deoxycholate Agar) media were prepared according to the manufacturers’ recommendations and pre-incubated at 37 °C for 24 h to verify sterility and microbiological quality. Each Salmonella spp. isolate was initially inoculated into 5 mL of BHI and incubated at 37 °C for 24 h. Subsequently, a 1 mL aliquot of the culture was transferred to 5 mL of Rappaport-Vassiliadis medium, with additional incubation for 24 h at 37 °C. After selective enrichment, aliquots of the cultures were seeded onto plates containing XLD medium. From the resulting growth, a single colony was selected, reinoculated into BHI, and incubated for an additional 24 h at 37 °C. Genomic DNA was extracted and purified using the PureLink™ kit (Thermo Fisher Scientific, United States), following the manufacturer’s instructions. Following enzymatic digestion with proteinase K, a modification was included in the protocol, consisting of an additional step of mechanical lysis via 10 consecutive pipetting movements (“up and down”) for each sample, with the aim of increasing the efficiency of bacterial genetic material release.

### Qualitative and quantitative DNA assessment

The integrity of the extracted DNA was assessed by 0.8% agarose gel electrophoresis with ethidium bromide, using 10 µL of each sample, based on the protocol by Sambrook and Russell [11], with minor adaptations. This step was used to verify the molecular integrity and apparent size of the extracted genomic DNA. DNA purity was assessed by spectrophotometry using a NanoDrop™ instrument (Thermo Fisher Scientific, United States), with 1.5 µL of each sample deposited directly onto the instrument’s sample well. The A260/280 and A260/230 ratios were considered as indicators of purity and possible contamination by proteins, phenols, or residual salts. Values close to 1.8–2.0 for A260/280 and 2.0–2.2 for A260/230 were considered adequate for subsequent steps [12]. Absolute DNA quantification was performed by fluorometry using a Qubit™ 4 fluorometer (Invitrogen, United States), with a specific kit for dsDNA. The concentrations obtained were used to adjust the samples prior to the preparation of the genomic libraries. After preparation, the libraries were normalized to 10 nM.

### Library preparation and whole-genome sequencing

Sequencing libraries were prepared using the Illumina DNA Prep kit (Illumina, USA) according to the manufacturer’s protocol. Sequencing was performed on the MiSeq i100 Plus platform (Illumina, USA) using a 2 × 150 bp paired-end strategy. Nine independent bacterial samples were processed, with a total yield compatible with an estimated minimum coverage of 30× per sample, sufficient for comparative analyses of bacterial genomes. The cartridge used was the MiSeq™ i100 Series Dry Cartridge 600 Cycles – 5M.

### Preprocessing of reads, reassembly, and genomic quality assessment

The raw reads obtained by sequencing underwent quality control using the fastp software, which removed adapters, trimmed bases with a quality score below Q30, and discarded reads shorter than 50 bp [13]. The reports generated in HTML and JSON formats were individually inspected for overall quality, GC content, duplication rate, and read size distribution. De novo assembly was performed using SPAdes v3.15 via the Shovill v1.1.0 pipeline, which integrates filtering, read organization, and optimized assembly for bacterial genomes [14]. Each sample generated two main files: spades.fasta, containing the raw contigs, and contigs.fa, containing the filtered contigs with a length ≥ 500 bp. Successful completion of the assembly was confirmed by the presence of the message “SPAdes finished” in the log files. The assembly graphs (contigs.gfa) were preserved for visual inspection in Bandage [15]. Assembly quality metrics were obtained using SeqKit v2.8.0, which was used to calculate the number of contigs, total genome size, longest contig, N50, L50, and GC content for each sample [16]. Comparison between the raw and filtered files showed that the removal of short contigs did not substantially alter the overall architecture of the assemblies, indicating the stability of the genome reconstruction process.

### Identification of the genetic determinants of resistance, virulence, biofilm formation, and persistence

The genotypic characterization of the isolates was based on the integration of multiple databases on resistance, virulence, and mobile genetic elements. The detection of antimicrobial resistance determinants was performed by integrating AMRFinderPlus, CARD, ResFinder, ARG-ANNOT, and MEGARes, thereby expanding the coverage of chromosomal and plasmid genes associated with resistance [17–21]. Virulence factors were identified based on the VFDB [22], while plasmid characterization was conducted using PlasmidFinder and supplemented by replicon reconstruction and typing with MOB-suite [23,24]. Extended screening was performed using AMRFinderPlus v3.12.8, with database version 2024-07-22.1, and with ABRicate against the CARD, ResFinder, ARG-ANNOT, MEGARes, VFDB, PlasmidFinder, Ecoh, Ecoli_VF, BacMet2, and NCBI databases. The analyses were processed with intensive use of parallelization. For disinfectant resistance genes, the BacMet2 database was searched using DIAMOND in blastp mode, since several relevant determinants were represented at the protein level. The alignments were subsequently converted to a format compatible with the final integration of the functional panel. As a complementary step, genes associated with biofilm formation and adhesion were retrieved by screening the VFDB and organized into biological modules, including curli (csgA–G), cellulose (bcsA, bcsB, bcsC, bcsZ, bcsE, bcsF, bcsG, bcsQ), type 1 fimbriae (fimA, fimC, fimD, fimF, fimG, fimH, fimZ, fimY), and global regulators such as csgD, adrA, yedQ, yddV, rpoS, flhDC, and fliC. The results from the different tools were consolidated using custom scripts, resulting in binary presence/absence matrices and curated functional panels.

### Sample selection for the structural phase and genotype–phenotype correlation

From the set of prioritized strains, a rational selection process was conducted for structural analysis and molecular docking. This selection considered three main criteria: i) the phenotypic profile of multidrug resistance previously observed in the antibiogram; ii) the presence of genes of biological interest related to antimicrobial resistance, efflux, and virulence; and iii) representativeness across serovars, years of isolation, and distinct biological profiles. The five selected strains, SH16.3, SH23.6, SM23.7, SM23.8, and SM23.9, proved suitable for this stage as they encompassed different combinations of phenotypic resistance, genetic repertoire, and epidemiological relevance. Subsequently, the protein sequences derived from these samples were compared with one another to remove redundancies. This procedure showed that some targets, such as AcrB and Sul2, were present with identical sequences in more than one isolate, allowing redundant proteins to be collapsed and the structural pipeline to be focused on unique targets. Thus, the final set of proteins subjected to structural analysis consisted of CTX-M-2 and SpvC, represented by SH16.3, and AcrB, CMY-2, and Sul2, represented by SH23.6. This strategy allowed us to rationally link the multidrug-resistant phenotype observed in the antibiogram with genetic determinants and their respective molecular targets, reducing pipeline redundancy without losing biological representativeness.

### Structural annotation, extraction of target proteins, and sequence validation

The genomes selected for the structural stage were annotated using Bakta, with a complete local database and multithreaded processing, to generate standardized annotation files in FAA, FFN, GFF3, GBFF, and TSV formats [25]. From the tabular and amino acid files, proteins annotated as β-lactamases, efflux transporters, folate metabolism enzymes, and virulence factors were retrieved. The extracted sequences of interest were compared for length and identity across the selected samples, allowing us to confirm redundancies and identify more representative variants. Next, for each structural target, the sequence derived from the genome was compared to the sequence present in the selected experimental structures in the Protein Data Bank (PDB), ensuring adequate coverage and alignment for docking.

### Selection and preparation of experimental three-dimensional structures

For targets with suitable experimental three-dimensional structures, the entries 1ZC2 (CMY-2), 7S2K (Sul2), 6Z12 (AcrB), and 4HAH (SpvC) were selected from the Protein Data Bank. The structures were chosen based on functional correspondence, sequence coverage, and suitability for molecular screening. The chains of interest were extracted using custom Python scripts, removing water molecules and non-essential heteroatoms. For Sul2, two versions of the receptor were prepared: one without cofactors and another preserving the Mg²⁺ ion present at the functional site of the 7S2K structure, in order to assess the impact of the metal on ligand binding. For CMY-2, AcrB, and SpvC, cleaned versions of the selected chain were used.

### Structural modeling of CTX-M-2

The CTX-M-2 target, derived from isolate SH16.3, underwent structural modeling using ColabFold v1.6.0 in a GPU-accelerated Linux environment, employing automatic templates, homology search via MMseqs2, and final relaxation using Amber [26]. The best model was selected based on the pipeline’s internal ranking. After modeling, the conserved catalytic motifs characteristic of class A β-lactamases were examined, including the SXXK motif, the SDN motif, and the KTG motif, which were used as a complementary check of the structural and functional consistency of the selected model. These residues were also used as a reference for positioning the docking box.

### Construction of chemical libraries and preparation of linkers

The ligand libraries were assembled in a target-directed manner and retrieved from PubChem in SDF format using a custom Python script [27]. For Sul2, classical sulfonamides and compounds related to the PABA/folate pathway were tested; for CMY-2 and CTX-M-2, a panel of β-lactams and β-lactamase inhibitors was used; for AcrB, classic substrates and inhibitors of efflux pumps were included; and for SpvC, an exploratory set of phosphorylated and phosphonated compounds. The molecules were organized into target-specific subsets, preserving pharmacological consistency between the biological mechanism and the chemical library employed.

### Molecular docking of reference compounds

Molecular docking was performed exclusively using GNINA, a program that integrates convolutional neural networks (CNNs) into the scoring and pose refinement process [28]. A single docking engine was used throughout the entire campaign to ensure the internal comparability of the results and avoid mixing different scoring functions. Scores were interpreted as relative metrics of structural prioritization, rather than as absolute estimates of experimental affinity [28,29]. For CMY-2 and Sul2, ligand-guided autoboxing using co-crystallized ligands was employed. For AcrB, the search was directed toward the distal pocket and the hydrophobic trap. For SpvC, the box was centered on the protein’s functional cavity. For CTX-M-2, the box was defined based on the catalytic residues identified in the structural model. For Sul2, the final docking run used for prioritization was performed against the 7S2K_A_keepMG.pdb receptor, with Mg²⁺ retained at the catalytic site. The search box was centered at x = 22.197; y = −1.245; z = 8.431, with dimensions of 22 × 22 × 22 Å, using --num_modes 10 and --exhaustiveness 16. For AcrB, docking was targeted at the distal pocket of the 6Z12_A_clean.pdb receptor, with the box centered at x = 92.023; y = 134.958; z = 131.903 and dimensions 29.500 × 24.903 × 28.782 Å, using -- num_modes 10 and --exhaustiveness 24. For CTX-M-2, the CTX-M-2_rank1_relaxed.pdb model was used, with a box defined from the catalytic residues 73, 76, 133–135, and 237–239, centered at x = 3.685; y = 5.648; z = −0.661 and dimensions 26.783 × 23.850 × 24.987 Å, with --num_modes 10 and --exhaustiveness 16. For SpvC, a large box was used around the functional region in 4HAH_A_clean.pdb, centered at x = −22.7345; y = −7.7895; z = 23.0470, with dimensions 58.741 × 56.901 × 51.664 Å, also with --num_modes 10 and --exhaustiveness 16. In all runs, poses were initially prioritized based on the Mode 1 score, supplemented, when necessary, by structural inspection of the recovered pose.

### Analog generation, cheminformatics-based prioritization, and redocking

Following the initial screening, a stage involving the generation of molecular analogs was carried out using RDKit, an open-source toolkit widely used in cheminformatics for reading and writing molecules, structural sanitization, fingerprint generation, descriptor calculation, and fragmentation and recombination operations [30]. For this step, the most promising parent compounds identified in each screening campaign were used as seed molecules. For the AcrB target, the seeds minocycline, doxycycline, tetracycline, tigecycline, and phenylalanine-arginine β-naphthylamide (PAβN) were employed. Analog generation was conducted using BRICS (Breaking of Retrosynthetically Interesting Chemical Substructures) rules, originally proposed for the decomposition and recombination of chemically plausible fragments in drug-like chemical spaces [31]. The parent molecules were read in SDF format, sanitized, and converted into canonical representations. Next, BRICS fragments were extracted and recombined with a maximum depth of 2, generating an expanded library of candidates. The resulting set was deduplicated using canonical SMILES, ensuring structural uniqueness in the final library. All generated analogs underwent 3D geometry construction in RDKit, through explicit hydrogen addition, conformational embedding, and geometric minimization using the UFF (Universal Force Field), which is useful for obtaining energetically plausible initial geometries in structural screening campaigns [32].

The cheminformatics-based prioritization of analogs was performed by calculating structural descriptors in RDKit, including molecular weight (MolWt), exact mass, cLogP, TPSA, number of hydrogen acceptors and donors, number of rotatable bonds, number of rings, number of heavy atoms, and formal charge. The QED (quantitative estimate of drug-likeness) index was also calculated, a metric proposed to rank compounds based on a trade-off between desirable physicochemical properties in bioactive molecules [33]. In addition, Morgan-type circular fingerprints were generated, and the Tanimoto similarity between each analog and the seed molecules was calculated.

The internal screening of the analogs was based on operational criteria defined for the campaign, requiring a molecular weight between 250 and 950 Da, cLogP between −1.5 and 7.5, TPSA between 40 and 260 Å², at least 2 rings, a maximum of 2 violations of drug-likeness rules, and a minimum similarity of 0.35 relative to the best seed molecule. Next, an internal PriorityScore was calculated by combining structural similarity, QED, penalty for violations, molecular weight proximity, and TPSA proximity, to rank the analogs and select candidates for redocking. The prioritized analogs were then redocked at the same molecular site used in the primary campaigns, preserving the receptor, box coordinates, number of modes, and exhaustiveness, in order to allow direct comparison between parent compounds and derivatives within a consistent protocol.

### Analysis of protein–ligand interactions and final prioritization of complexes

After docking, the best complexes were analyzed using PLIP (Protein–Ligand Interaction Profiler), a tool developed for the automated detection of hydrogen bonds, hydrophobic contacts, π-stacking, cation–π interactions, salt bridges, halogen bonds, metal–ligand interactions, and water bridges in protein–ligand complexes [34]. For each prioritized ligand, the best structural conformation was selected to characterize noncovalent interactions and identify recurring residues in the molecular recognition of the pocket. The final prioritization of the complexes was based on the integration of docking score, pose consistency, interaction recovery by PLIP, and the compounds’ cheminformatics profile. This strategy allowed us to distinguish candidates with numerically favorable scores but low interactional support from those that combined good relative energy, a coherent interaction network, and a more plausible physicochemical profile, thereby rationally guiding the selection of complexes for subsequent stages of structural refinement and molecular dynamics.

### Additional analyses and supplementary material

Circular genome maps are provided as Supplementary Figures S1 and S2, and additional cheminformatics and structural panels are provided as Supplementary Figures S3-S5. In the supplementary material, the selected genomes were represented as circular maps in Proksee, based on the assembled contigs and corresponding annotation files, highlighting the distribution of resistance and virulence genes and other genomic features of interest [35]. Similarly, phylogenetic trees and comparative representations of the isolates were included as a complementary analysis, integrating the genomic context and molecular prioritization into a unified analytical strategy.

## RESULTS

### Phenotypic profile and selection rationale for the strains

The five Salmonella enterica isolates included in the study had previously been classified as multidrug-resistant based on antibiotic susceptibility testing, as they exhibited resistance to three or more classes of antimicrobials. This result provided the experimental starting point for subsequent genomic and structural investigations, as it indicated the presence of potentially multiple resistance mechanisms, combining enzymatic determinants, efflux systems, regulatory alterations, and possibly mobile elements associated with horizontal gene acquisition. Among the observed phenotypic profiles, SH16.3 exhibited resistance to gentamicin, enrofloxacin, nalidixic acid, and tetracycline, while SH23.6 showed resistance to colistin, amoxicillin/clavulanic acid, and enrofloxacin.

In contrast, S. Minnesota isolates exhibited broader resistance profiles: SM23.7 was resistant to ampicillin, chloramphenicol, cefotaxime, tetracycline, enrofloxacin, and sulfamethoxazole-trimethoprim; SM23.8 was resistant to ampicillin, tetracycline, enrofloxacin, and sulfamethoxazole-trimethoprim, as well as additional resistance to amoxicillin/clavulanic acid and ceftiofur in a supplementary assay; and SM23.9 showed resistance to ampicillin, aztreonam, cefotaxime, tetracycline, azithromycin, and sulfamethoxazole-trimethoprim, as well as additional resistance to amoxicillin/clavulanic acid. These profiles supported the selection of strains for the downstream pipeline, as they encompassed different combinations of resistance to β-lactams, sulfonamides, tetracyclines, quinolones/fluoroquinolones, phenicols, and polymyxins. The phenotypic profile supporting strain selection is summarized in Figure 2. Thus, the experimental rationale of the study was supported by the convergence between phenotypic data and the genetic profile, reinforcing that the strategy for prioritizing molecular targets was aligned with the experimental behavior of the strains.

**Figure 2.**
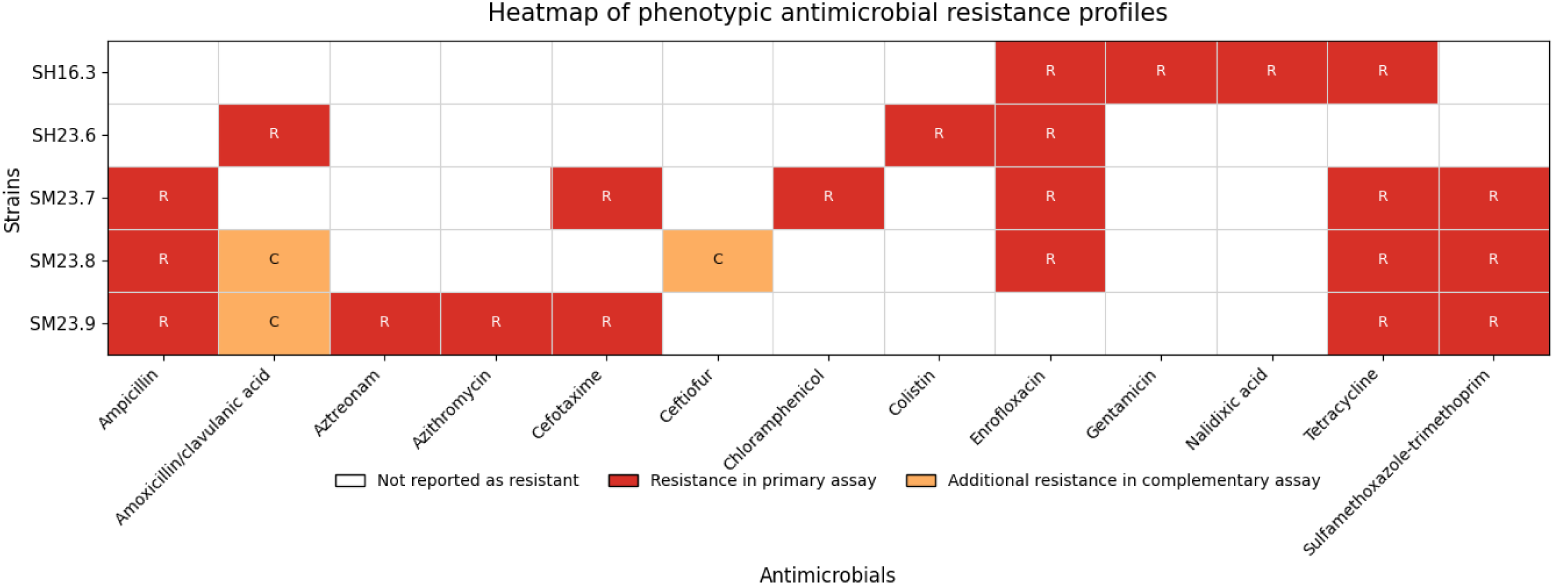
Phenotypic antimicrobial resistance profiles of the five prioritized Salmonella enterica isolates. Heatmap summarizing the antimicrobial resistance profiles used to support isolate selection and downstream genomic and structure-based analyses.

### Sequencing, genome assembly, and assembly quality

Paired-end sequencing generated sufficient coverage for the reconstruction of the five bacterial genomes and yielded high-quality assemblies for most of the samples. In four of the five genomes analyzed, the total reconstructed size ranged from approximately 4.78 to 4.95 Mb, with a GC content of around 52.1%, values consistent with Salmonella enterica. The assemblies also exhibited satisfactory continuity, with N50 values ranging from approximately 225 kb to 440 kb and a relatively low number of contigs, facilitating gene annotation and subsequent comparative analyses. In contrast, sample SH16.3 behaved as a striking outlier in the dataset. This assembly had a size of 6.42 Mb, 4,432 contigs, an N50 of only 4,160 bp, and a largest contig of 83,346 bp, in addition to a slight deviation in GC content, at 51.79%. This pattern suggests a combination of greater fragmentation, a potentially high plasmid load, the presence of repetitive regions or an expanded mobilome, as well as greater structural complexity compared to the other strains. Despite this, the priority targets of biological interest were clearly identified in the annotation of this sample, which allowed it to be included in the downstream pipeline. Thus, although SH16.3 exhibited inferior assembly quality compared to the other strains, this limitation did not preclude the extraction of specific target genes. On the contrary, its unique profile proved particularly relevant for the identification of exclusive targets, such as CTX-M-2 and SpvC.

### Resistome, virulome, and consistency with the observed phenotype

Functional annotation and targeted screening revealed a resistance pattern that was biologically consistent with the previously observed multidrug-resistant antibiotic susceptibility profile. Among the five genomes analyzed, two main profiles stood out. The first was represented by strain SH16.3, in which blaCTX-M-2, sul1, tet(A), qacE/qacEdelta1, and the virulence factor spvC were identified. This set is consistent with a complex phenotype, involving resistance to extended-spectrum β-lactams, sulfonamides, and tetracyclines, as well as possible increased tolerance to quaternary ammonium compounds. The simultaneous presence of blaCTX-M-2 and spvC was particularly relevant, as it brought together in a single isolate a clinical resistance determinant and a virulence effector associated with bacterial survival and modulation of host-microbe interaction. The second profile was observed mainly in strains SH23.6, SM23.7, SM23.8, and SM23.9, in which the recurrent combination of acrB, sul2, and tet(A) was detected, accompanied, in part of the group, by genes from the CMY family. In SH23.6 and SM23.7, the presence of blaCMY-2 was clear and structurally consistent; in SM23.9, a member of the CMY family was detected, though with a reduced length compared to canonical CMY-2; whereas in SM23.8, the profile was dominated by AcrB, Sul2, and TetA, with no equally clear hit for CMY-2 in the regions prioritized for modeling. This pattern reinforces the idea that the resistance observed experimentally did not depend on a single mechanism, but rather on a functional arrangement combining enzymatic inactivation, active drug efflux, and classic determinants of acquired resistance. In interpretive terms, the association between tet(A), sul1/sul2, and β-lactamase genes offers a qualitatively consistent explanation for the low susceptibility to tetracyclines, sulfonamides, and β-lactams observed in multidrug-resistant strains. Concurrently, the presence of acrB, ramA, acrA, acrR, and other components of efflux systems suggests an additional layer of tolerance to multiple compounds, consistent with broader phenotypic profiles and persistence under antimicrobial pressure. Regarding virulence and persistence, the functional matrix also showed significant conservation of genes related to curli, biofilm, and adhesion, especially among S. Minnesota isolates. Genes such as csgA–G and csgD were widely detected, indicating maintenance of the central apparatus for extracellular matrix formation and colonization. The distribution of the main resistance, efflux, biofilm, and virulence-associated genes is shown in Figure 3. Taken together, the results suggest that the isolates studied combine antimicrobial resistance with the potential for environmental persistence and colonization, a characteristic particularly relevant in the contexts of animal production and the food chain.

**Figure 3.**
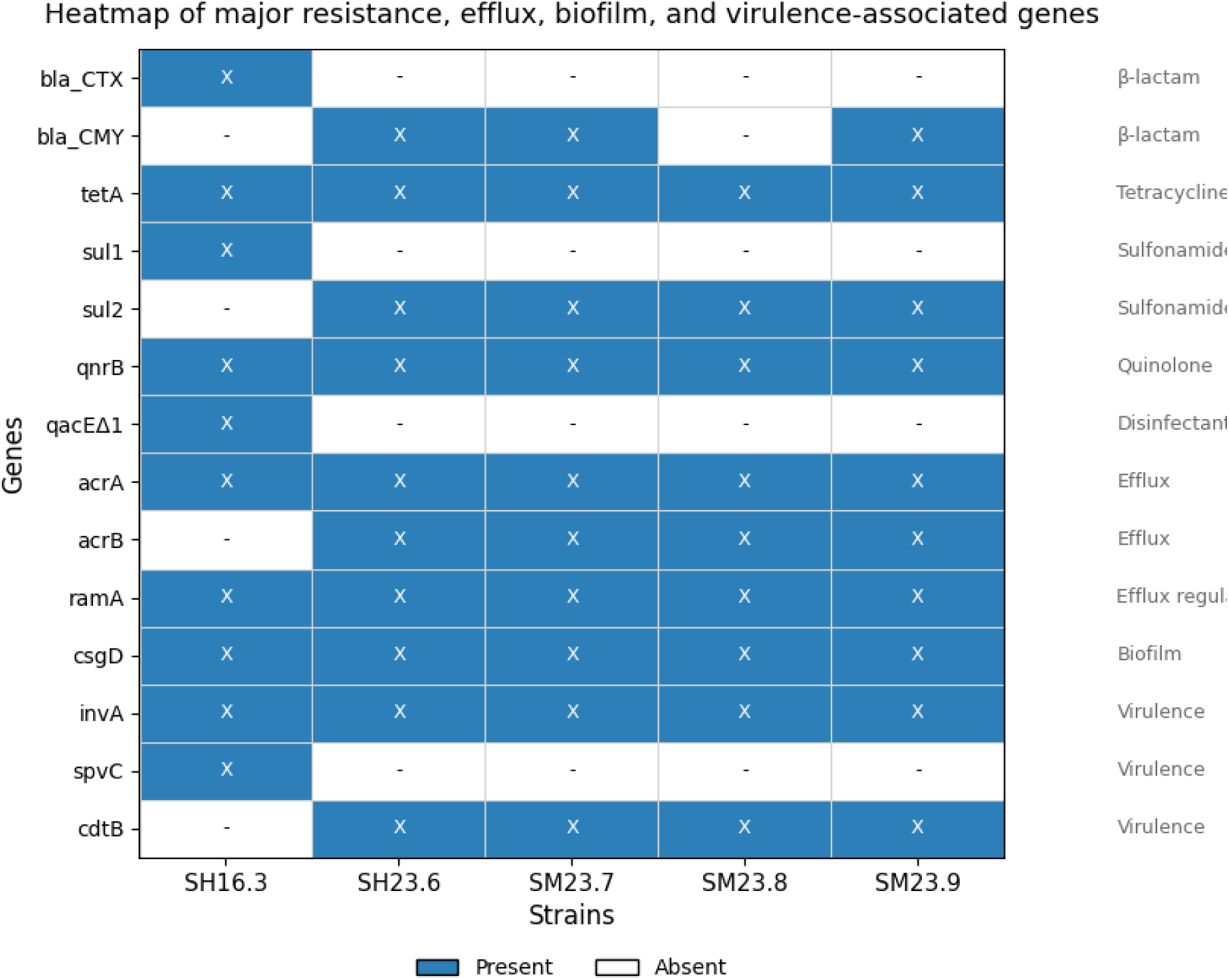
Heatmap of major resistance, efflux, biofilm, and virulence-associated genes. Blue cells indicate presence and white cells indicate absence of the corresponding gene in each Salmonella enterica strain selected for structural analysis.

### Prioritization of molecular targets for the structural phase

The integration of genomics, biological relevance, and structural availability led to the prioritization of five key targets for the docking and molecular optimization phase: CTX-M-2, SpvC, AcrB, CMY-2, and Sul2. These targets represented different mechanistic classes: a class A β-lactamase, a virulence effector, an RND efflux pump, a class C β-lactamase, and a sulfonamide-resistant dihydropteroate synthase. The extracted protein sequences confirmed the robustness of this selection. AcrB was 1,049 amino acids long and was identical among SH23.6, SM23.7, SM23.8, and SM23.9, indicating a highly conserved structural target among the recent isolates analyzed. The same pattern was observed for Sul2, with 271 amino acids and an identical sequence among these four strains. For CMY-2, the targets from SH23.6 and SM23.7 exhibited 381 amino acids and complete identity, whereas SM23.9 presented a shorter variant of the CMY family. SpvC and CTX-M-2, exclusive to the unique profile of SH16.3, had lengths of 241 and 291 amino acids, respectively. These findings were important for two reasons. First, they showed that the structural domain was not fragmented into multiple, poorly comparable variants, but anchored in conserved and biologically relevant targets. Second, they justified the use of a reduced number of representative structures without loss of biological coverage within the analyzed set.

### Correspondence between the target sequence and experimental structures

The validation between the target sequence and the three-dimensional structure was one of the strongest points of the pipeline. The alignments between the selected bacterial proteins and the structures available in the PDB showed virtually perfect correspondence. For CMY-2, the comparison with the 1ZC2 structure revealed 100.0% identity and 100.0% coverage. For Sul2, the 7S2K structure also showed 100.0% identity and full coverage. The same was true for SpvC with the 4HAH structure, again with 100.0% identity and full coverage. For AcrB, structure 6Z12 showed 99.81% identity and full coverage, with only two substitutions across 1,049 residues. This result had a direct methodological implication: the CMY-2, Sul2, SpvC, and AcrB targets could be docked using experimental structures that were practically isogenic to the actual bacterial proteins. In terms of structural robustness, this scenario is clearly superior to the use of distantly related homologous models and lends a high degree of confidence to the geometry of the analyzed sites.

### Structural modeling of CTX-M-2 and validation of the predictive model

Since no experimental structure was directly incorporated into the pipeline for the CTX-M-2 target of strain SH16.3, a predictive structure was built using ColabFold/AlphaFold2. The best model obtained was the rank_001 relaxed model, with a pLDDT of 93.7 and a pTM of 0.883, values consistent with a model of high global reliability. In addition to the global metrics, sequence inspection confirmed the presence of the catalytic motifs expected in class A β-lactamases: the SXXK motif at position 73, the SDN motif at 133–135, and the KTG motif at 237–239. The simultaneous preservation of these three functional elements reinforced the model’s suitability for catalytic site-oriented docking. Thus, unlike the other targets, CTX-M-2 was docked against a predicted receptor, but with consistent evidence of structural reliability and functional conservation.

### Molecular docking of reference compounds Sul2

The docking campaign for Sul2 revealed a methodologically significant result already in the comparison between receptors with and without the crystallographic metal. When docking was performed without retaining Mg²⁺, some of the compounds exhibited inconsistent or unfavorable scores, including positive values for molecules that should be compatible with the functional site. In contrast, retaining the metal in the receptor (keepMG) produced a more coherent ranking from a chemical and structural standpoint. In this scenario, sulfamethoxazole was the best compound in the primary set, with −7.50 kcal/mol, followed by p-aminobenzoic acid (−7.18 kcal/mol) and sulfametizole, in the expanded set, with −7.12 kcal/mol. Other sulfonamide analogs showed intermediate or poor performance, including sulfadoxine, sulfametazine, sulfisoxazole, sulfatiazol, dapsone, and sulfadiazine. Taken together, the results suggest that the Sul2 site was sensitive to the metal-dependent context and that the structural representation with the cofactor preserved was the most realistic option for the campaign.

### CTX-M-2

Docking against the CTX-M-2 model yielded an even more pronounced profile for broad-spectrum cephalosporins. Ceftriaxone was the best ligand, with −8.61 kcal/mol, followed by cefepime (−8.08 kcal/mol) and cefotaxime (−8.07 kcal/mol). Next came ampicillin (−7.70 kcal/mol), aztreonam (−7.60 kcal/mol), and avibactam (−7.19 kcal/mol). Compounds such as amoxicillin, ceftazidime, clavulanic acid, tazobactam, and sulbactam performed less well. This result was particularly interesting because it reinforced the functional relevance of CTX-M-2 as a target associated with extended-spectrum β-lactam resistance in the SH16.3 isolate. Although docking does not replace kinetic assays, the observed pattern was consistent with strong structural recognition of cephalosporins by the modeled catalytic site.

### SpvC

For the virulence effector SpvC, the best results were obtained with phosphorylated or phosphonated compounds that are structurally compatible with the type of chemistry recognized by this protein. The best score was observed for O-phospho-L-tyrosine (−6.25 kcal/mol), followed by phenylphosphonic acid (−5.31 kcal/mol) and 4-nitrophenyl phosphate (−5.22 kcal/mol). Other compounds in the set, such as O-phospho-L-threonine, O-phospho-L-serine, glyphosate, fosfomycin, phosphonoacetic acid, and phosphonoformate, exhibited less favorable scores. These data support the plausibility of the chosen site and indicate that the target preferentially responds to ligands containing phosphorylated groups or phosphorous mimetics, consistent with its functional nature.

### AcrB

The AcrB target exhibited one of the most promising profiles in the primary screening. In the distal pocket, the best compound in the reference set was phenylalanine-arginine β-naphthylamide (PAβN), with −8.67 kcal/mol, followed by doxycycline (−8.50 kcal/mol) and tetracycline (−7.80 kcal/mol). Tigecycline showed −7.19 kcal/mol, while minocycline, chloramphenicol, ciprofloxacin, florfenicol, norfloxacin, and nalidixic acid exhibited less favorable values. The target-specific docking rankings of the reference compounds are summarized in Figure 4. This ranking was important for two reasons. First, it showed that the distal pocket defined for docking was capable of recognizing molecules known to be associated with transport, efflux, or functional modulation of the system. Second, it provided a solid baseline for the optimization step using analogs, allowing direct comparison between the parent compounds and the derivatives generated in RDKit.

**Figure 4.**
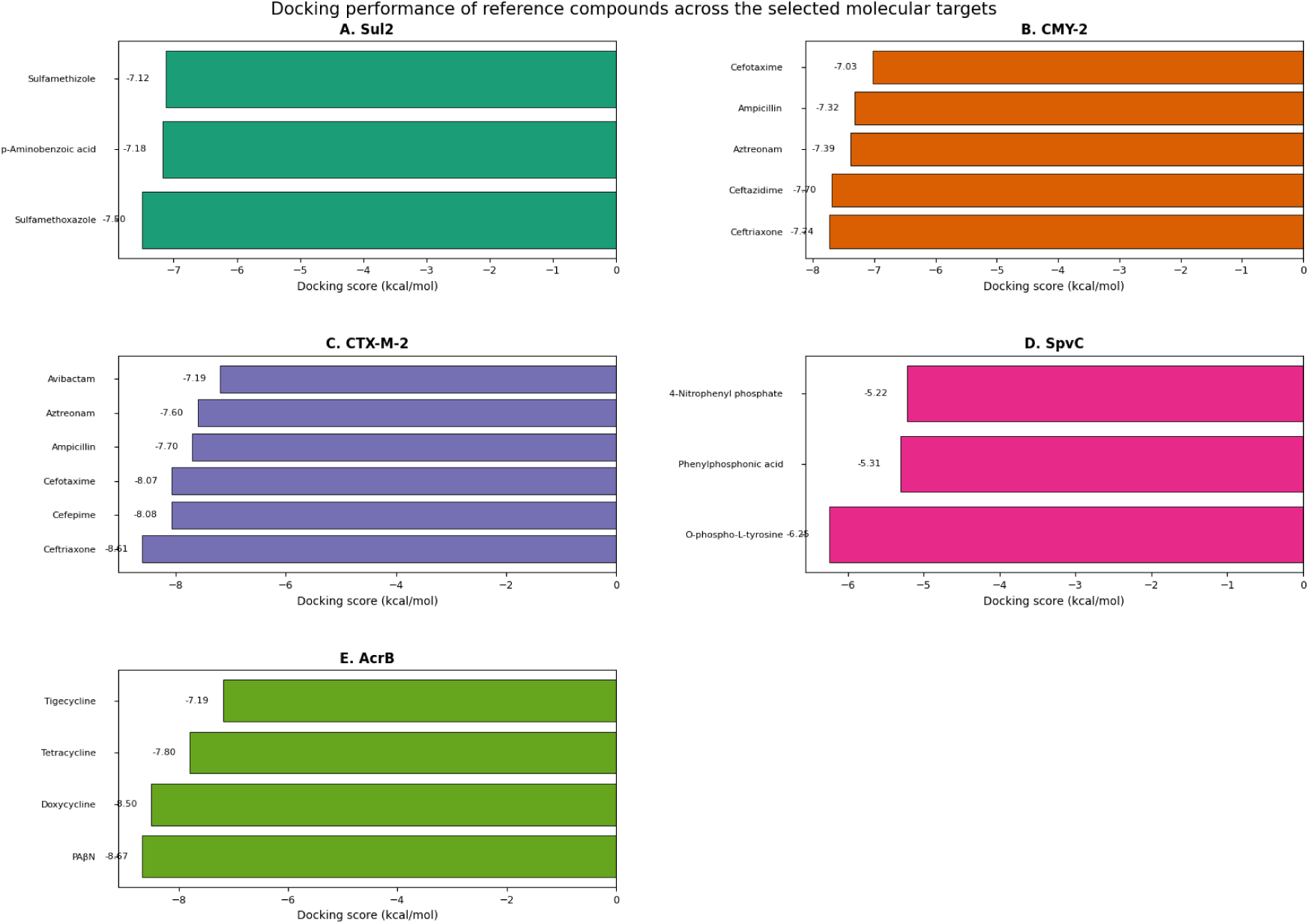
Docking performance of reference compounds across the selected molecular targets. Target-specific docking rankings obtained with GNINA for the reference ligand libraries, providing the baseline for subsequent analog generation and redocking.

### Lead optimization: comparative performance in Sul2 and AcrB

#### Sul2

The library of Sul2 analogs generated by RDKit yielded a reduced set of valid candidates, of which 95 passed internal filters after prioritization. The best compounds ranked by cheminformatics profile included sulfametazine, sulfametizole, sulfisoxazole, sulfamethoxazole, and the derivative BRICS_00001. However, when these compounds were redocked on the same Sul2 receptor, the structural gain was limited.

The best scores were −7.50 kcal/mol for sulfametizole and −7.44 kcal/mol for sulfamethoxazole, followed by sulfisoxazole (−7.02 kcal/mol). The new derivative BRICS_00001 had a substantially lower score (−4.43 kcal/mol), and sulfametazine performed poorly (−3.36 kcal/mol). Taken together, these results indicate that the optimization step on Sul2 did not yield derivatives clearly superior to the best parent compounds. Thus, although the pipeline was methodologically sound, the target’s structural response was relatively conservative and did not support a strong expansion of the lead optimization program for this protein.

#### AcrB

The situation was different for AcrB. The library of analogs generated from the most relevant parent compounds yielded 234 valid molecules, of which 95 remained after filtering. Among the prioritized compounds, several analogs had higher scores than the reference parent compounds, indicating a real structural gain in the distal pocket. The best results observed were ACRB_BRICS_00197 (−10.99 kcal/mol), ACRB_BRICS_00124 (−10.80 kcal/mol), ACRB_BRICS_00089 (−10.33 kcal/mol), followed by ACRB_BRICS_00076, ACRB_BRICS_00115, and ACRB_BRICS_00206, all at −9.65 kcal/mol. In purely energetic terms, these values clearly surpassed the best parental compounds, including PAβN (−8.67 kcal/mol) and doxycycline (−8.50 kcal/mol), suggesting that the exploration of chemical space via BRICS was more successful in AcrB than in Sul2. However, the final interpretation was not based solely on the score. The PLIP analysis showed that the hits with the best structural support were ACRB_BRICS_00124, ACRB_BRICS_00076, ACRB_BRICS_00206, and ACRB_BRICS_00115. Among them, ACRB_BRICS_00124 emerged as the most consistent candidate, with −10.80 kcal/mol, 6 hydrogen bonds, 3 hydrophobic contacts, and 1 salt bridge. ACRB_BRICS_00076 exhibited −9.65 kcal/mol, with 5 hydrogen bonds and 8 hydrophobic contacts, indicating strong accommodation in the aromatic and hydrophobic regions of the pocket. ACRB_BRICS_00206 showed a balanced profile, with 4 hydrogen bonds, 4 hydrophobic contacts, and 1 π-stacking interaction, while ACRB_BRICS_00115 exhibited a favorable but more moderate profile, with 3 hydrogen bonds, 3 hydrophobic contacts, and 1 π-stacking. Recurring residues observed among the best complexes included Asn81, Gln89, Ser133, Ser134, Phe615, Phe617, Glu673, and Asn719, suggesting consistent occupation of the same functional region of the distal pocket of AcrB. This pattern reinforces the idea that the analogs not only numerically improved the scores but also preserved a plausible structural recognition logic at the target site.

On the other hand, two compounds require a more cautious interpretation. Although ACRB_BRICS_00197 and ACRB_BRICS_00089 achieved the highest scores in the set, PLIP did not identify any explicit non-covalent interactions for these complexes. This does not automatically invalidate these candidates, but it does indicate that they should not be treated as priority hits. The AcrB analog optimization and interaction-supported prioritization are shown in Figure 5A-D. Thus, from an integrated perspective of score and interaction network, ACRB_BRICS_00124 should be considered the best supported hit in the set, followed by ACRB_BRICS_00076 and ACRB_BRICS_00206.

**Figure 5A.**
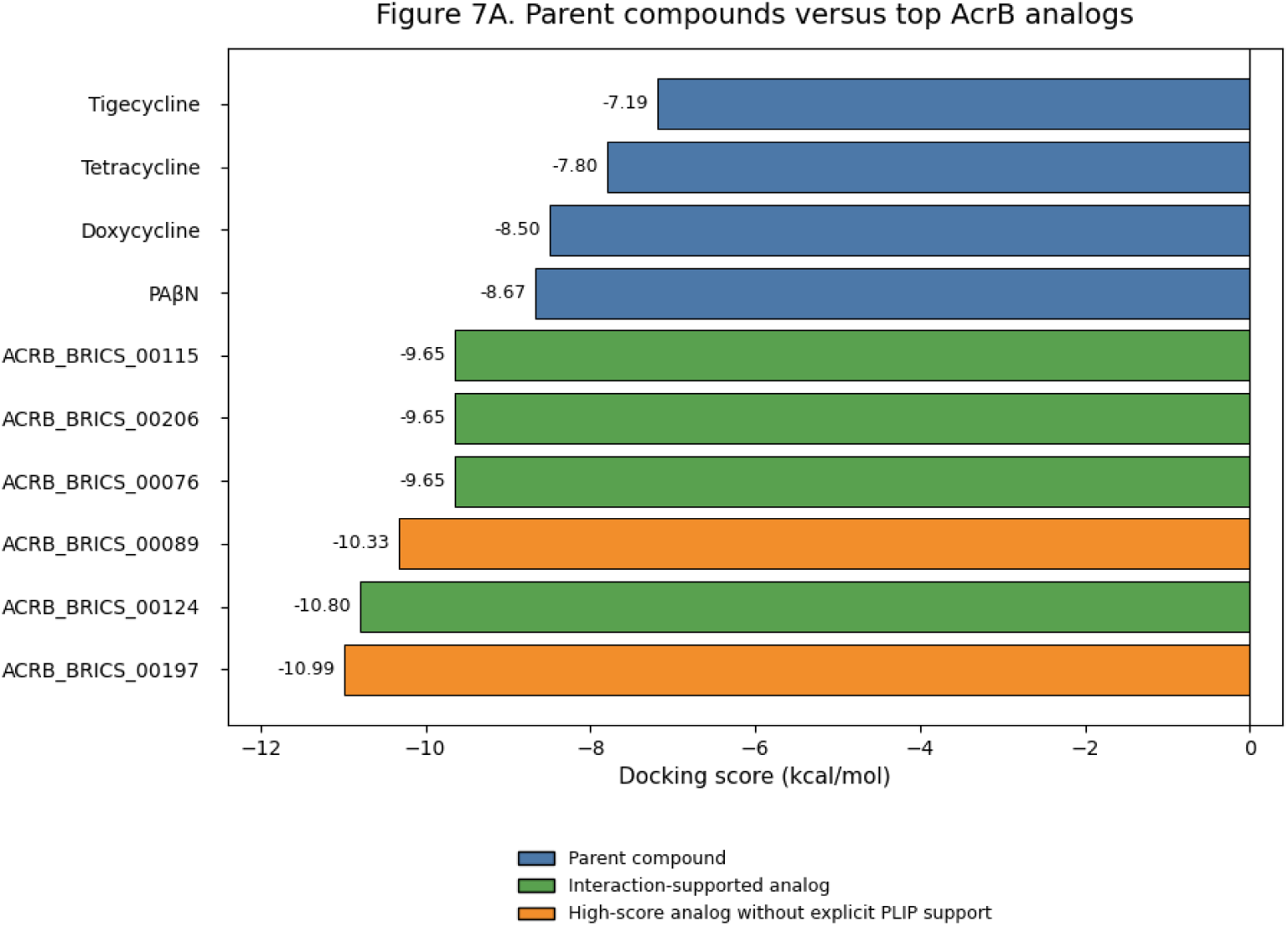
Parent compounds versus top AcrB BRICS-derived analogs. Comparison between reference parent compounds and top BRICS-derived analogs docked against the AcrB distal pocket. Green bars indicate analogs supported by both docking score and PLIP interaction profiles, orange bars indicate high-scoring analogs lacking explicit PLIP interaction recovery, and blue bars represent parental reference compounds.

**Figure 5B.**
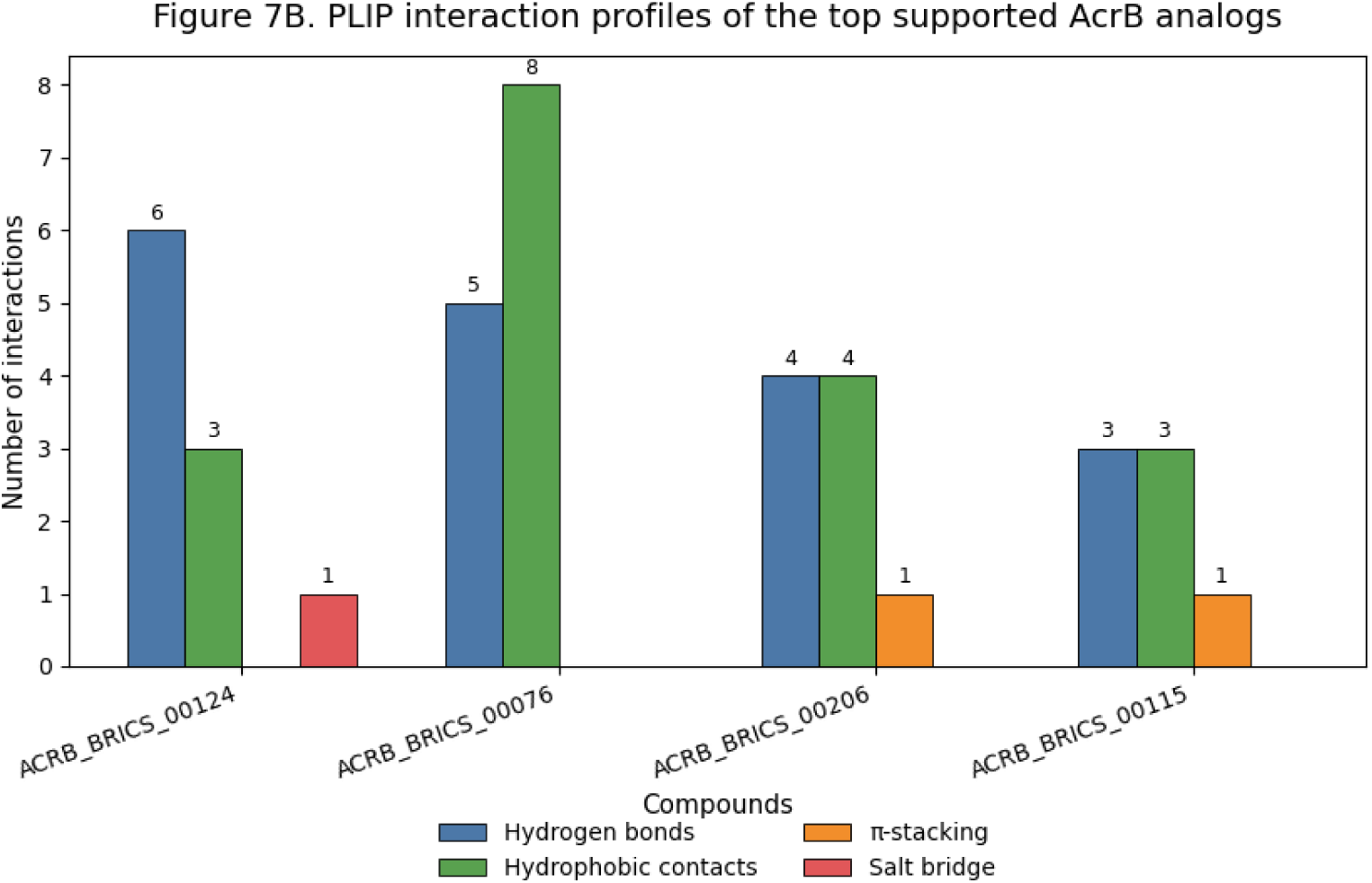
PLIP interaction profiles of the top supported AcrB analogs. Interaction summary for the main AcrB analogs prioritized after integration of docking score and non-covalent interaction recovery.

**Figure 5C.**
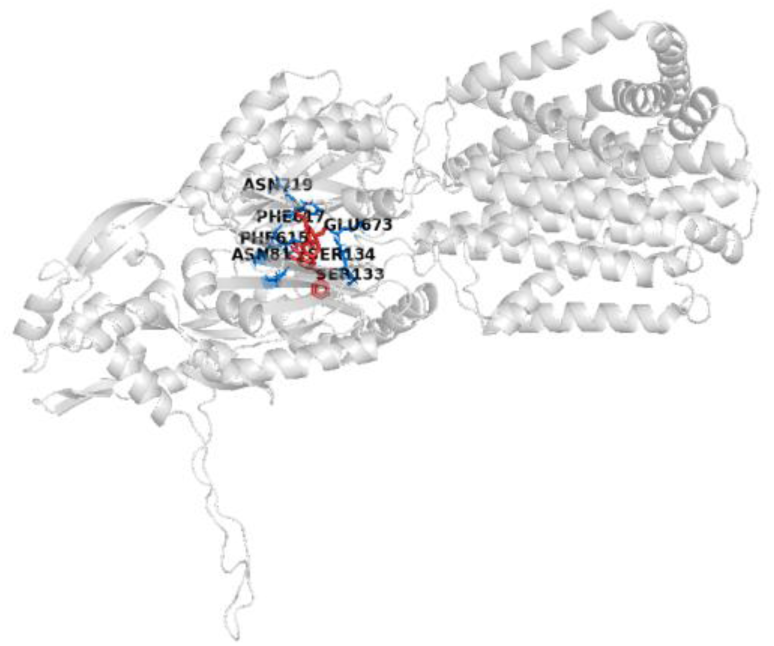
Docking pose of ACRB_BRICS_00124 in the AcrB distal pocket. Close-up representation of the predicted binding pose and the surrounding residues involved in molecular recognition.

**Figure 5D.**
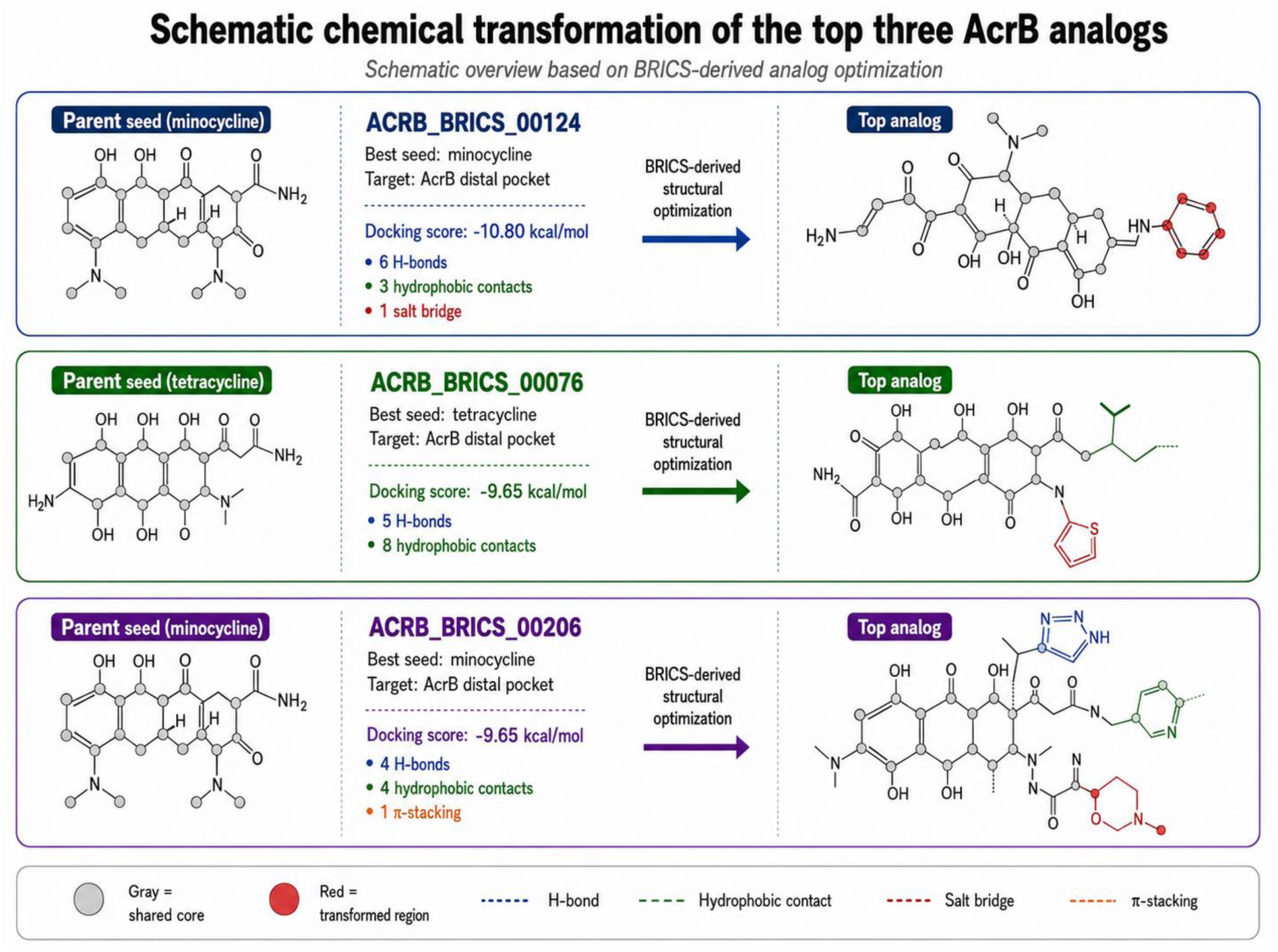
Schematic chemical transformation of the top three interaction-supported AcrB analogs. The final supported hits were ACRB_BRICS_00124 (−10.80 kcal/mol), ACRB_BRICS_00076 (−9.65 kcal/mol), and ACRB_BRICS_00206 (−9.65 kcal/mol). ACRB_BRICS_00124 and ACRB_BRICS_00206 were associated with minocycline-derived analog generation, whereas ACRB_BRICS_00076 was associated with tetracycline-derived analog generation.

### Integration of phenotype, genomics, and molecular prioritization

When analyzed together, the results indicate overall consistency between the experimental behavior of the strains, the detected genetic content, and the structural prioritization of targets. The multidrug susceptibility testing provided the phenotypic background. Genomics revealed biologically plausible combinations of genes associated with resistance to β-lactams, sulfonamides, and tetracyclines, as well as efflux systems and persistence genes. Sequence–structure validation showed that the prioritized targets were anchored to experimental receptors virtually identical to the actual bacterial proteins or, in the case of CTX-M-2, to a highly reliable model.

At the functional level, the results also allowed us to distinguish two behaviors. For Sul2, docking and the analog generation step supported the target as biologically relevant, but chemical optimization did not convincingly outperform the parent compounds. For AcrB, however, the analog generation and selection step produced consistent improvements in binding scores and interactions, establishing this target as the most promising avenue in the rational ligand optimization program conducted in this study.

This difference between the targets is, in itself, an important finding. Rather than forcing a uniform narrative for all proteins, the data indicate that the chemical space explored was more responsive in the distal pocket of AcrB, whereas Sul2 exhibited more restrictive structural behavior. Consequently, the results obtained justify prioritizing the best AcrB complexes for the next stages of structural refinement, including molecular dynamics and a more in-depth evaluation of the conformational stability of protein–ligand recognition.

## DISCUSSION

The results of this study reinforce the notion that the emergence and persistence of Salmonella Heidelberg and Salmonella Minnesota in the Brazilian poultry industry should not be interpreted solely as epidemiological phenomena related to serovar frequency, but rather as the expression of strains adapted to intensive production environments, frequently associated with multidrug resistance and genetic profiles compatible with persistence and dissemination. On a broader scale, genomic analyses have demonstrated that S. Heidelberg and S. Minnesota are among the most relevant serovars in chickens produced in Brazil and in Brazilian poultry products detected abroad, suggesting adaptive success and potential public health impact beyond the local context [2].

The convergence between phenotype and genotype was one of the strongest findings of the study. The detection of blaCTX-M-2, blaCMY-2, sul1/sul2, tet(A), and components of the AcrAB-TolC system provides a mechanistically plausible explanation for the resistance profiles observed in the antibiogram, especially against β-lactams, sulfonamides, tetracyclines, and fluoroquinolones. This pattern is consistent with Brazilian studies that identified, in S. Heidelberg and S. Minnesota of avian origin, the predominance of blaCMY-2, frequently associated with IncC plasmids, as well as combinations of genes that support multidrug phenotypes and increase the potential for horizontal spread of resistance [3]. Converging data from national surveys have shown high and increasing rates of resistance among Salmonella isolates from chicken meat in Brazil, particularly among the Heidelberg and Minnesota serovars [36].

The contrast between sample SH16.3 and the other isolates is also noteworthy. Although this strain exhibited a more fragmented assembly and a larger total genome than expected for the group, the priority targets were clearly recovered, allowing it to remain in the analytical pipeline. Rather than invalidating the sample, this behavior suggests greater structural complexity, possibly related to an expanded mobilome, repetitive regions, plasmid content, or genomic rearrangements. This interpretation is biologically plausible for multidrug-resistant Salmonella, especially when ESBL genes and accessory determinants may be associated with mobile elements. The concurrent presence of blaCTX-M-2 and spvC in SH16.3 is particularly interesting, as it brings together, in a single isolate, a clinically relevant resistance marker and a virulence effector associated with the modulation of host MAPK pathways. Although the present study did not functionally evaluate SpvC activity, the literature shows that this protein acts as a phosphothreonine lyase and contributes to the attenuation of the host inflammatory response [37,38].

Another strength of the study was the sequence-structure validation. For CMY-2, Sul2, SpvC, and AcrB, the near-perfect correspondence between the proteins extracted from the isolates and the experimental structures selected from the PDB substantially reduces one of the main sources of uncertainty in docking studies: the use of receptors that are structurally distant from the actual sequence of the target organism. This aspect strengthens the internal interpretation of the screening campaigns, as it brings the computational system closer to the molecular reality of the studied isolates. For CTX-M-2, although it was necessary to employ a predictive model, the high pLDDT and pTM values, combined with the preservation of the classic catalytic motifs of class A β-lactamases, support the model’s suitability for initial exploration of the active site. Nevertheless, in accordance with the standards of rigor expected by high-impact journals, it is important to recognize that even high-quality models do not replace experimental structural validation, especially when subtle differences in geometry can influence ligand pose and binding affinity.

In the case of Sul2, the results highlight a methodologically relevant observation: maintaining Mg²⁺ at the functional site produced more consistent rankings than the version without the cofactor. This result suggests that the receptor’s metal-dependent environment directly influences ligand binding and recognition, which is consistent with the fact that small changes in the site representation can significantly alter docking scores. Although sulfamethoxazole and some analogs exhibited relatively favorable performance, the chemical expansion stage did not yield derivatives clearly superior to the parent compounds. In interpretive terms, this suggests that the chemical space explored for Sul2 was more restrictive, or that the initial set of fragments and seeds was not sufficiently diverse to break the recognition pattern already established by the target. Thus, the data support Sul2 as a biologically relevant target, but do not indicate, at this stage, a strong axis of structural optimization comparable to that observed for AcrB.

AcrB has emerged as the most promising target of the study, from both biological and cheminformatics perspectives. This interpretation is supported by three converging lines of evidence. First, AcrB was detected in multiple isolates with a highly conserved sequence, indicating that the target represents a stable trait of the bacterial population analyzed. Second, the distal pocket/hydrophobic trap selected for docking is recognized in the literature as a critical region for the development of RND pump inhibitors, including the interaction of compounds in this phenylalanine-rich cavity [39,40]. Third, the BRICS-based analog generation step produced compounds with a significant increase in score relative to the parent compounds, and some of these hits also showed plausible interaction support via PLIP. In particular, ACRB_BRICS_00124, ACRB_BRICS_00076, and ACRB_BRICS_00206 combined favorable scores with coherent networks of hydrogen bonds, hydrophobic contacts, and, in some cases, π interactions, suggesting consistent occupation of the prioritized functional region. In a broader context, these findings are consistent with the renewed interest in efflux inhibitors for Gram-negative bacteria, as RND pumps remain among the primary drivers of multidrug resistance and continue to be targets of high strategic value [9,10,40].

Nevertheless, the interpretation of docking results should remain deliberately conservative. Docking scores, even in neural network-enhanced engines such as GNINA, do not equate to experimental affinities and may overestimate certain poses when the receptor representation, conformational flexibility, solvation, or system dynamics are not fully captured. Recent studies emphasize that deep learning-based docking and scoring methods expand screening capabilities but still exhibit significant limitations regarding generalization, calibration, and experimental translation. Thus, the fact that two compounds with excellent scores in AcrB did not produce explicit interactions in PLIP is a pertinent methodological caveat, not a contradiction. In practice, this means that the energy ranking should be interpreted in conjunction with the pose, chemical consistency, recoverability of interactions, and, ideally, subsequent refinement steps, such as molecular dynamics and phenotypic validation.

From a translational perspective, the study also clearly illustrates the current role of WGS as a platform for functional prioritization, rather than merely descriptive surveillance. Recent reviews highlight that whole-genome sequencing has revolutionized outbreak investigation and the characterization of foodborne pathogens by integrating typing, traceability, and genetic profiles of resistance and virulence. In this sense, the strategy adopted here—starting from the phenotype, integrating genomics, reducing protein redundancy, validating the structural suitability of targets, and only then proceeding to molecular screening—represents a robust analytical framework well aligned with current trends in applied genomic microbiology [7]. The fact that different targets responded differently to chemical screening is also relevant, as it shows that the integration of molecular epidemiology and cheminformatics can identify, even at an early stage, which pathways warrant greater experimental investment.

In summary, the data obtained support three main conclusions. First, the analyzed isolates fit into the contemporary landscape of multidrug-resistant poultry Salmonella in Brazil, especially regarding the importance of S. Heidelberg and S. Minnesota as reservoirs of resistance genes and persistence traits [2,4–6]. Second, the consistency between the antibiogram, genetic content, and structural prioritization confirms the utility of an integrative approach for the rational selection of targets. Third, among the proteins investigated, AcrB stood out as the target most responsive to chemoinformatic optimization, while Sul2 exhibited more conservative behavior, and CTX-M-2 and SpvC remained biologically relevant targets, though still dependent on further experimental and structural investigation. Thus, the central contribution of this study is not to claim the discovery of ready-to-use inhibitors, but to demonstrate, based on consistent genomic and structural evidence, a rational pathway for prioritizing targets and molecular candidates in multidrug-resistant avian strains of Salmonella.

## CONCLUSION

The present study demonstrated that the analyzed poultry isolates of Salmonella enterica, belonging to the S. Heidelberg and S. Minnesota serovars, exhibit a consistent pattern of multidrug resistance, supported by the convergence of phenotypic, genomic, and structural data. The integration of antibiotic susceptibility testing, whole-genome sequencing, and functional screening enabled the identification of biologically plausible combinations of genes associated with resistance to β-lactams, sulfonamides, tetracyclines, and other antimicrobials, as well as determinants linked to efflux, persistence, and virulence. These findings reinforce the epidemiological relevance of these serotypes in the Brazilian poultry industry and demonstrate that the approach employed was capable of transforming surveillance data into a rational strategy for molecular prioritization. Among the targets evaluated, AcrB stood out as the most promising axis for chemoinformatics exploration, as it exhibited conservation among isolates, biological consistency with the multidrug-resistant phenotype, and a favorable response during the analog generation and redocking stage, with candidates that combined higher scores and a plausible interaction network. In contrast, Sul2 proved to be biologically relevant but less responsive to structural optimization, while CTX-M-2 and SpvC remained targets of interest, primarily due to their functional and epidemiological value, although they require further experimental investigation. Thus, the study did not merely. Among the targets evaluated,

AcrB stood out as the most promising target for cheminformatics exploration, as it exhibited conservation across isolates, biological consistency with the multidrug-resistant phenotype, and a favorable response during the analog generation and redocking stages, with candidates that combined higher scores and a plausible interaction network. In contrast, Sul2 proved to be biologically relevant but less responsive to structural optimization, while CTX-M-2 and SpvC remained targets of interest, primarily due to their functional and epidemiological value, although further experimental investigation is required. Thus, the study not only genetically characterizes multidrug-resistant Salmonella strains of poultry importance but also establishes an integrative and reproducible pipeline for target selection and initial ligand prioritization, providing a solid foundation for future investigations involving structural refinement, molecular dynamics, and biological validation of the most promising complexes

## SUPPLEMENTARY FIGURES

**Supplementary Figure S1.**
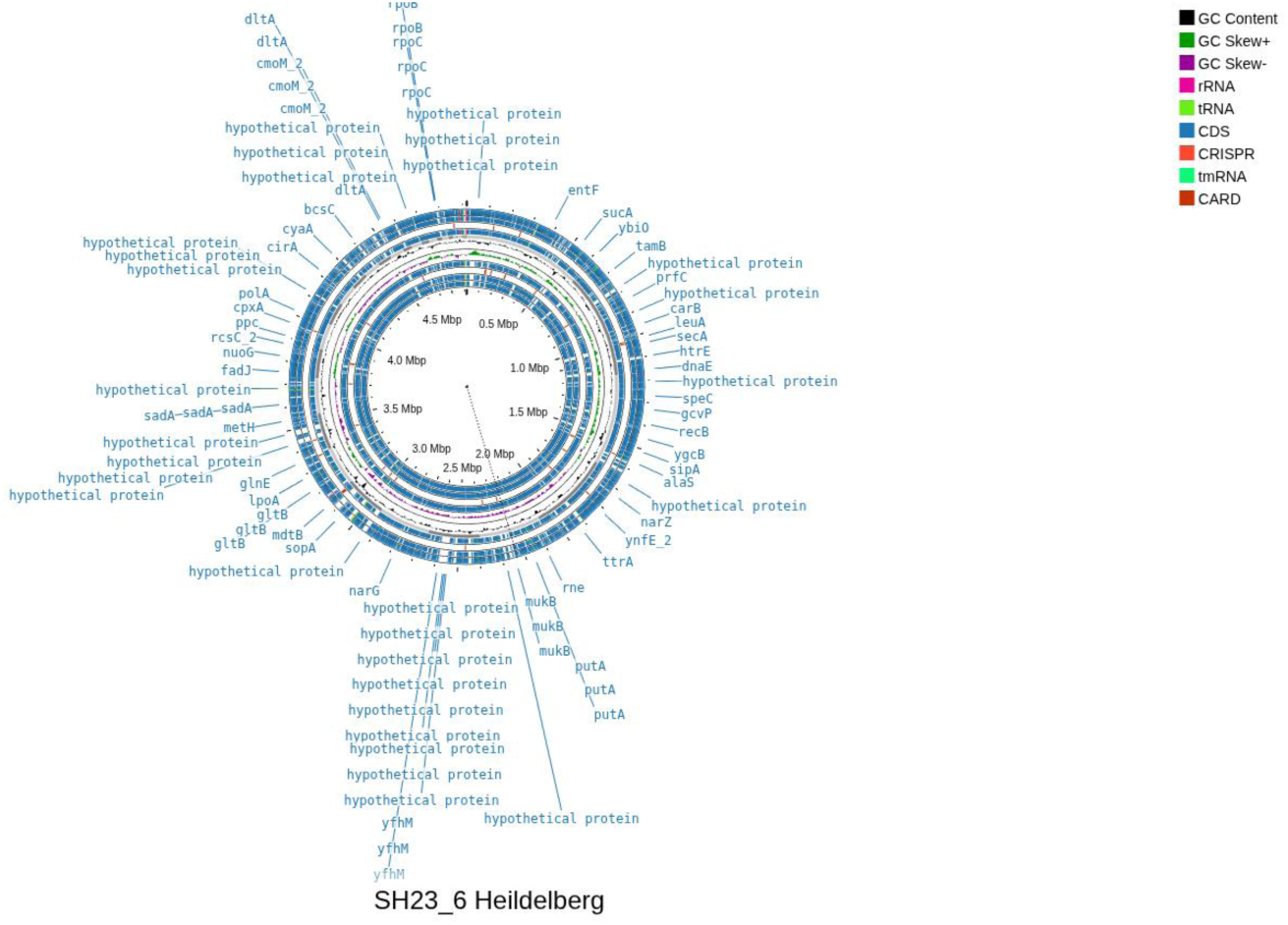
Circular genome map of Salmonella enterica serovar Heidelberg strain SH23.6. Proksee-generated representation showing coding sequences on both strands, RNA features, antimicrobial resistance determinants, CRISPR/Cas-associated elements, GC content, and GC skew.

**Supplementary Figure S2.**
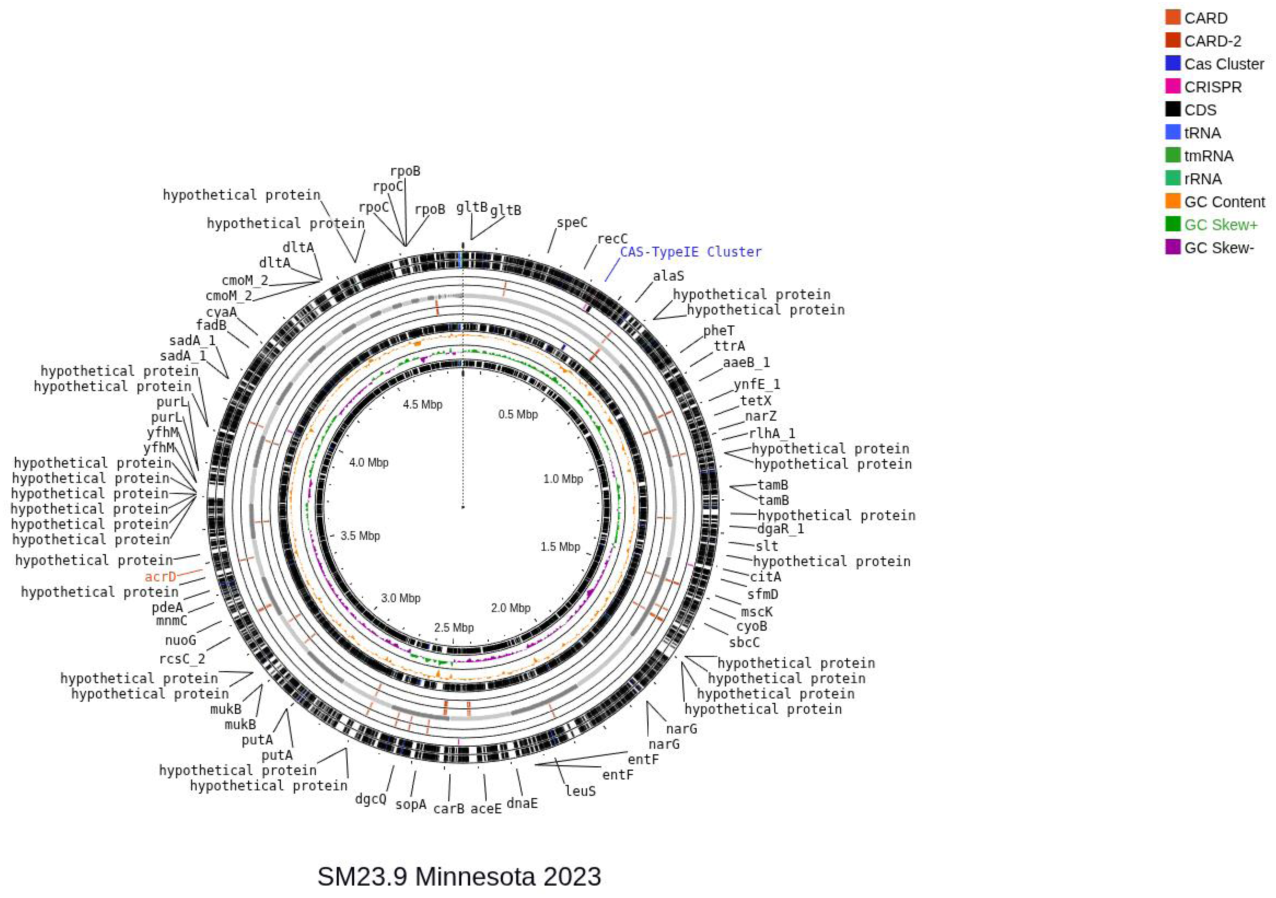
Circular genome map of Salmonella enterica serovar Minnesota strain SM23.9. The outer rings show annotated coding sequences and RNA features, whereas internal rings summarize antimicrobial resistance determinants, CRISPR/Cas-associated features, GC content, and GC skew.

**Supplementary Figure S3.**
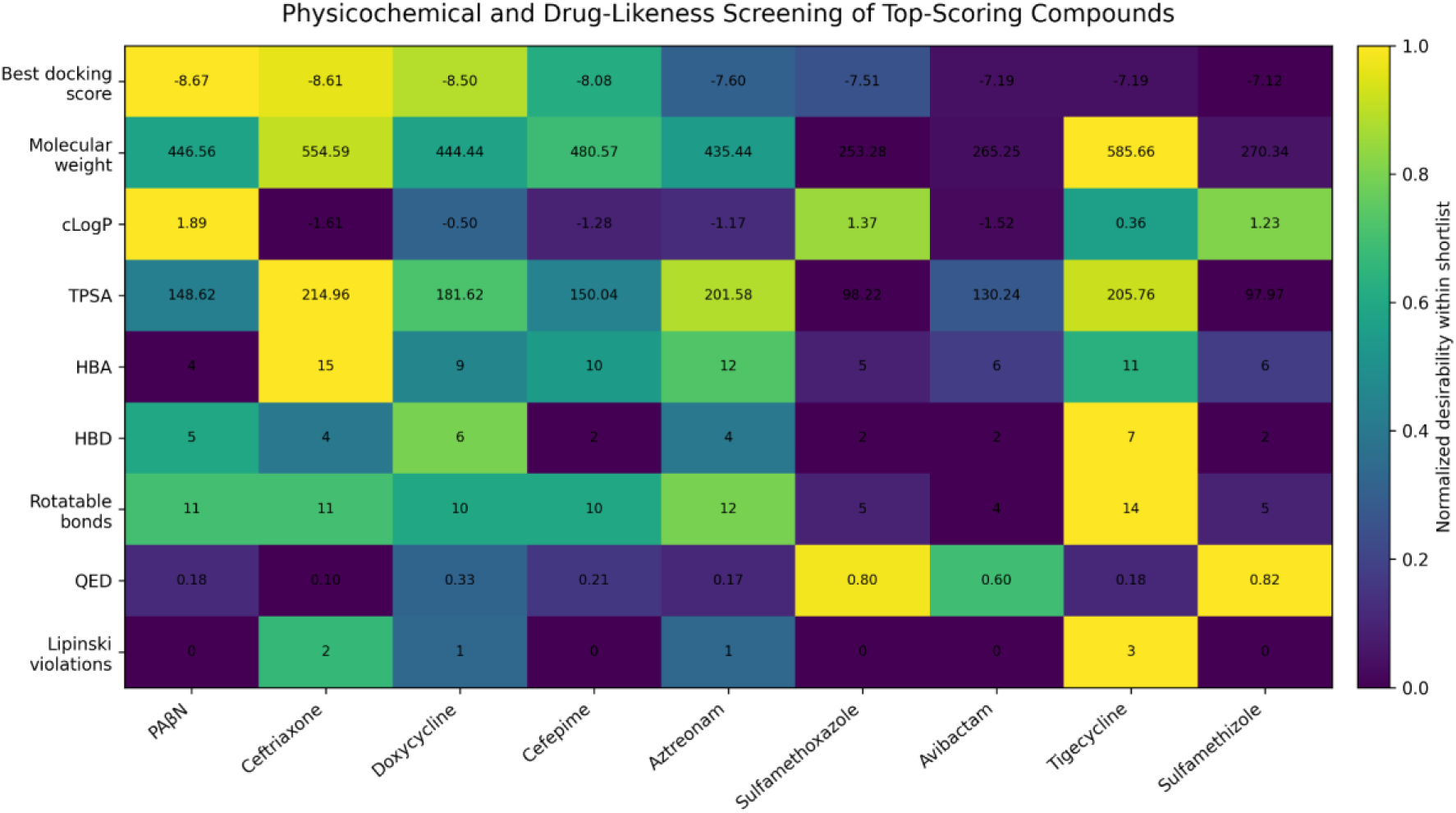
Physicochemical and drug-likeness screening of top-scoring compounds. Heatmap summarizing selected molecular descriptors used to compare and prioritize compounds during the cheminformatics stage.

**Supplementary Figure S4.**
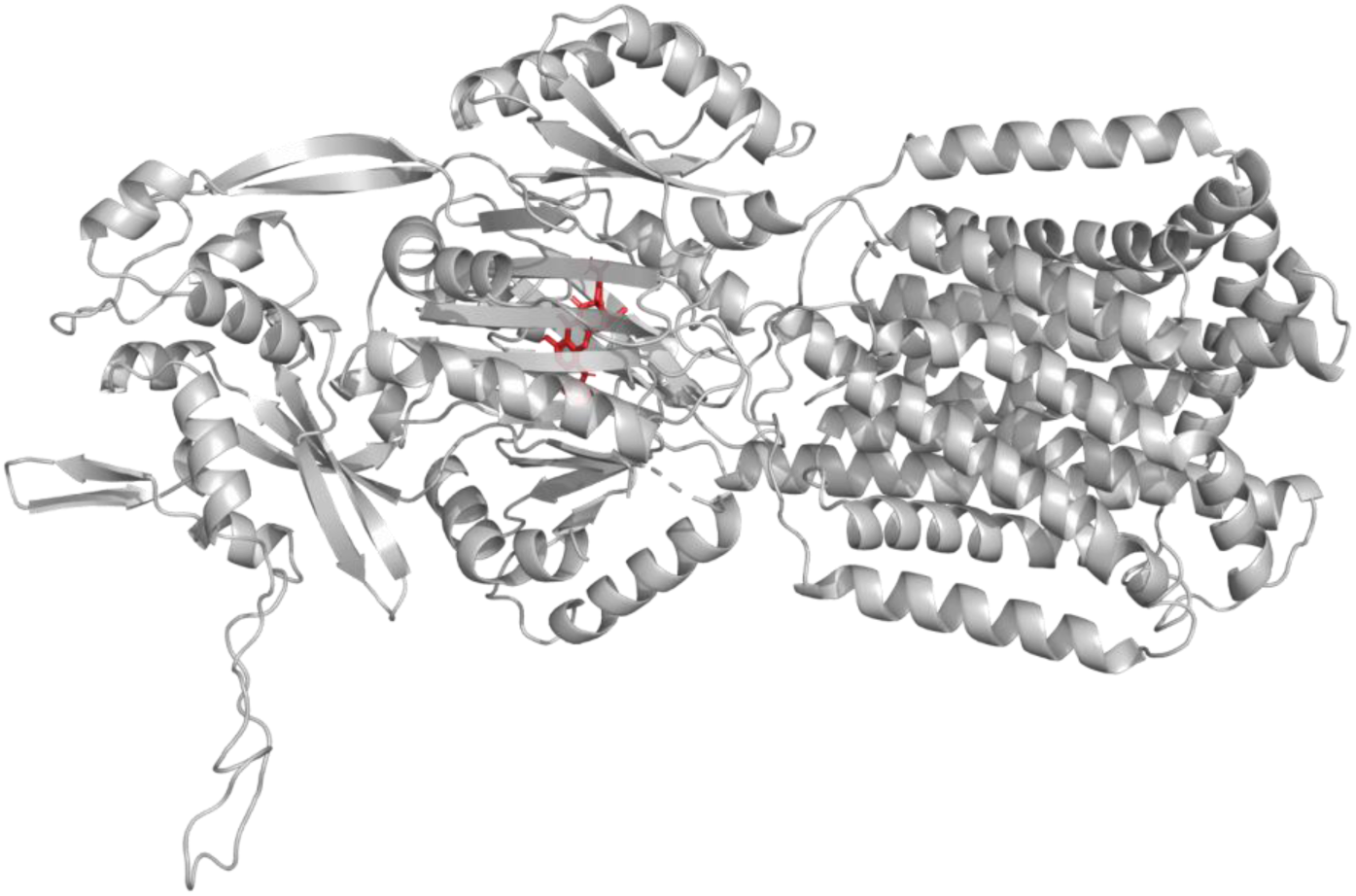
Overview of the AcrB docking complex. Global structural view of the AcrB receptor with the prioritized ligand pose highlighted as supplementary support for the close-up view shown in Figure 5C.

**Supplementary Figure S5.**
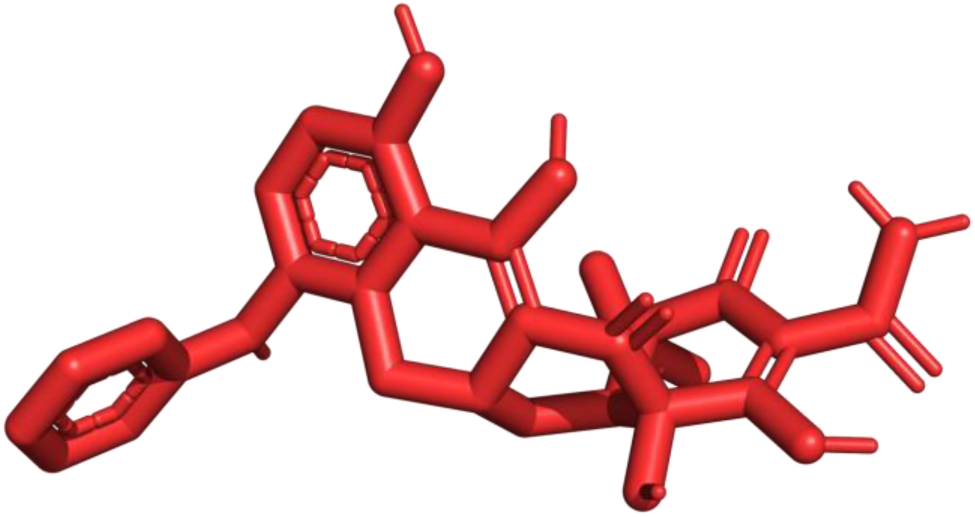
Isolated three-dimensional representation of the prioritized AcrB analog used for structural visualization.

## Notes

### Competing Interest Statement

The authors have declared no competing interest.

## REFERENCES

1. World Health Organization. Salmonella (non-typhoidal). Geneva: WHO; 2018. Available from: WHO institutional portal. Accessed 2026 Mar 29.

2. Alikhan NF, et al. Dynamics of Salmonella enterica and antimicrobial resistance in the Brazilian poultry industry and global impacts on public health. PLoS Genet. 2022;18(6):e1010174. doi:10.1371/journal.pgen.1010174.

3. Saidenberg ABS, et al. Salmonella Heidelberg and Salmonella Minnesota in Brazilian broilers: genomic characterization of third-generation cephalosporin and fluoroquinolone-resistant strains. Environ Microbiol Rep. 2023;15(1):119–128. doi:10.1111/1758-2229.13132.

4. De Almeida Figueira A, et al. Phenotypic and genomic assessment of antimicrobial resistance and virulence factors determinants in Salmonella Heidelberg isolated from broiler chickens. Animals. 2025;15(7):1003. doi:10.3390/ani15071003.

5. Mesa D, et al. An insight into the characterization of multidrug resistant Salmonella Minnesota isolated from poultry in Brazil. Infect Genet Evol. 2026;139:105896. doi:10.1016/j.meegid.2026.105896.

6. Justino L, et al. Environmental persistence and genotypic and phenotypic characterization of Salmonella Minnesota in poultry slaughterhouses. Pathogens. 2026;15(3):247. doi:10.3390/pathogens15030247.

7. Gomes E, et al. Advances in whole genome sequencing for foodborne pathogens: implications for clinical infectious disease surveillance and public health. Front Cell Infect Microbiol. 2025;15:1593219. doi:10.3389/fcimb.2025.1593219.

8. European Food Safety Authority. Whole genome sequencing in foodborne outbreaks. EFSA institutional portal. Accessed 2026 Mar 30.

9. El-Demerdash AS, et al. Natural inhibitors of Salmonella MDR efflux pumps AcrAB and AcrD: an integrated in silico, molecular, and in vitro investigation. Int J Mol Sci. 2024;25(23):12949. doi:10.3390/ijms252312949.

10. Schuster S, et al. Comparative reassessment of AcrB efflux inhibitors reveals differential impact of specific pump mutations on the activity of potent compounds. Microbiol Spectr. 2024;12(2):e03045–23. doi:10.1128/spectrum.03045-23.

11. Sambrook J, Russell DW. Molecular cloning: a laboratory manual. 3rd ed. Cold Spring Harbor: Cold Spring Harbor Laboratory Press; 2001.

12. Thermo Fisher Scientific. Assessment of nucleic acid purity. Wilmington: Thermo Fisher Scientific; 2015. Technical Note TN52646-E-0215M.

13. Chen S, et al. fastp: an ultra-fast all-in-one FASTQ preprocessor. Bioinformatics. 2018;34(17):i884–i890. doi:10.1093/bioinformatics/bty560.

14. Bankevich A, et al. SPAdes: a new genome assembly algorithm and its applications to single-cell sequencing. J Comput Biol. 2012;19(5):455–477. doi:10.1089/cmb.2012.0021.

15. Wick RR, et al. Bandage: interactive visualization of de novo genome assemblies. Bioinformatics. 2015;31(20):3350–3352. doi:10.1093/bioinformatics/btv383.

16. Shen W, et al. SeqKit: a cross-platform and ultrafast toolkit for FASTA/Q file manipulation. PLoS One. 2016;11(10):e0163962. doi:10.1371/journal.pone.0163962.

17. Feldgarden M, et al. Validating the AMRFinder tool and resistance gene database by using antimicrobial resistance genotype-phenotype correlations in a collection of isolates. Antimicrob Agents Chemother. 2019;63(11):e00483–19. doi:10.1128/AAC.00483-19.

18. Jia B, et al. CARD 2017: expansion and model-centric curation of the comprehensive antibiotic resistance database. Nucleic Acids Res. 2017;45(D1):D566–D573. doi:10.1093/nar/gkw1004.

19. Zankari E, et al. Identification of acquired antimicrobial resistance genes. J Antimicrob Chemother. 2012;67(11):2640–2644. doi:10.1093/jac/dks261.

20. Gupta SK, et al. ARG-ANNOT, a new bioinformatic tool to discover antibiotic resistance genes in bacterial genomes. Antimicrob Agents Chemother. 2014;58(1):212–220. doi:10.1128/AAC.01310-13.

21. Lakin SM, et al. MEGARes: an antimicrobial resistance database for high throughput sequencing. Nucleic Acids Res. 2017;45(D1):D574–D580. doi:10.1093/nar/gkw1009.

22. Chen L, et al. VFDB: a reference database for bacterial virulence factors. Nucleic Acids Res. 2005;33(Database issue):D325–D328. doi:10.1093/nar/gki008.

23. Carattoli A, et al. In silico detection and typing of plasmids using PlasmidFinder and plasmid multilocus sequence typing. Antimicrob Agents Chemother. 2014;58(7):3895–3903. doi:10.1128/AAC.02412-14.

24. Robertson J, Nash JHE. MOB-suite: software tools for clustering, reconstruction and typing of plasmids from draft assemblies. Microb Genom. 2018;4(8):e000206. doi:10.1099/mgen.0.000206.

25. Schwengers O, et al. Bakta: rapid and standardized annotation of bacterial genomes via alignment-free sequence identification. Microb Genom. 2021;7(11):000685. doi:10.1099/mgen.0.000685.

26. Mirdita M, et al. ColabFold: making protein folding accessible to all. Nat Methods. 2022;19(6):679–682. doi:10.1038/s41592-022-01488-1.

27. Kim S, et al. PubChem 2025 update. Nucleic Acids Res. 2025;53(D1):D1516–D1525. doi:10.1093/nar/gkae1059.

28. McNutt AT, et al. GNINA 1.0: molecular docking with deep learning. J Cheminform. 2021;13:43. doi:10.1186/s13321-021-00522-2.

29. Trott O, Olson AJ. AutoDock Vina: improving the speed and accuracy of docking with a new scoring function, efficient optimization, and multithreading. J Comput Chem. 2010;31(2):455–461.

30. RDKit. RDKit: open-source cheminformatics. Release 2024_09_2. Zenodo; 2024.

31. Degen J, Wegscheid-Gerlach C, Zaliani A, Rarey M. On the art of compiling and using drug-like chemical fragment spaces. ChemMedChem. 2008;3(10):1503–1507.

32. Rappe AK, et al. UFF, a full periodic table force field for molecular mechanics and molecular dynamics simulations. J Am Chem Soc. 1992;114(25):10024–10035.

33. Bickerton GR, et al. Quantifying the chemical beauty of drugs. Nat Chem. 2012;4(2):90–98.

34. Salentin S, et al. PLIP: fully automated protein-ligand interaction profiler. Nucleic Acids Res. 2015;43(W1):W443–W447.

35. Grant JR, et al. Proksee: in-depth characterization and visualization of bacterial genomes. Nucleic Acids Res. 2023;51(W1):W484–W492. doi:10.1093/nar/gkad326.

36. Rau RB, et al. Antimicrobial resistance of Salmonella from poultry meat in Brazil: results of a nationwide survey. Epidemiol Infect. 2021;149:e126. doi:10.1017/S0950268821001110.

37. Mazurkiewicz P, et al. SpvC is a Salmonella effector with phosphothreonine lyase activity on host mitogen-activated protein kinases. Mol Microbiol. 2008;67(6):1371–1383. doi:10.1111/j.1365-2958.2008.06134.x.

38. Li H, et al. The phosphothreonine lyase activity of a bacterial type III effector family. Science. 2007;315(5814):1000–1003. doi:10.1126/science.1138960.

39. Aron Z, Opperman TJ. The hydrophobic trap: the Achilles heel of RND efflux pumps. Res Microbiol. 2018;169(7-8):393–400. doi:10.1016/j.resmic.2017.11.001.

40. Duffey M, et al. Inhibitors of Gram-negative bacterial efflux pumps. ACS Infect Dis. 2024;10(5):1458–1482. doi:10.1021/acsinfecdis.4c00091.

